# A mycobacterial Sec61 inhibitor disrupts lysosome function by blocking Vacuolar-ATPase biosynthesis

**DOI:** 10.1101/2025.08.26.671788

**Authors:** Belinda S Hall, Kwabena Owusu-Boateng, Rachel E Simmonds

**Affiliations:** Discipline of Microbes, Infection and Immunity, School of Bioscience and Medicine, Faculty of Health and Medical Science, University of Surrey, Guildford, UK

**Keywords:** lysosome, Vacuolar-ATPase, mycolactone, Sec61

## Abstract

Mycolactone is the virulence toxin of *Mycobacterium ulcerans*, causative agent of Buruli ulcer. Mycolactone inhibits the Sec61-dependent co-translational translocation of signal peptide-bearing secreted and membrane proteins into the endoplasmic reticulum. Sec61 inhibition leads to accumulation of mislocalised proteins in the cytosol and initially triggers an integrated stress response-dependent activation of autophagy that contributes to cell survival. Here we show that sustained exposure of cells to mycolactone causes a block in late-stage autophagy and induces nuclear translocation of the lysosomal stress marker TFEB. This coincides with loss of ATP6AP1 and ATP6AP2, Sec61-substrates required for assembly of the Vacuolar-ATPase, leading to reduced lysosomal biogenesis and acidification. This compromises the cell’s capacity to withstand the proteostatic stress caused by Sec61 inhibition and impairs the ability of phagocytes to combat infection with *M. ulcerans*. Loss of lysosomal function could contribute to both the tissue necrosis and immunosuppression seen in Buruli ulcer. Furthermore, since Sec61 inhibition is being pursued as a therapeutic target in several diseases, potential drugs should be screened against this activity to avoid unwanted side-effects.

## Introduction

Buruli ulcer is a chronic skin infection caused by *Mycobacterium ulcerans*. Lesions can display extensive tissue necrosis but are painless with little inflammation. Mycolactone, a polyketide lactone, is the sole identified virulence factor synthesised by *M. ulcerans* and is responsible for all the pathology of the disease (George *et al*, 1999; Yotsu *et al*, 2018). Mycolactone has cytotoxic, immunosuppressive and analgesic properties *in vitro* and *in vivo* (Baron *et al*, 2016; George *et al*, 2000; Guenin-Mace *et al*, 2011; Marion *et al*, 2014). The main cellular target of mycolactone is the Sec61 translocon, an oligomeric complex which translocates nascent polypeptides into the endoplasmic reticulum (Bhadra *et al*, 2021; Hall *et al*, 2014). Mycolactone binds directly to Sec61α, the pore-forming component of the complex, blocking access of signal-peptide bearing proteins to the channel, and preventing opening of the plug domain that keeps the resting translocon closed (Gerard *et al*, 2020; Itskanov *et al*, 2023). Translocation of secreted proteins, Type I and II single-pass transmembrane proteins, and a subset of multi-pass membrane proteins with signal peptides or long internal loops is inhibited (Hsieh *et al*, 2025; McKenna *et al*, 2017).

The inhibition of Sec61 by mycolactone explains the immunosuppression seen in Buruli ulcer, since immune cells can no longer produce the surface receptors and cytokines needed for both innate and acquired immune responses. It is also the driver of mycolactone’s cytotoxicity, since cells expressing Sec61α mutants that no longer bind mycolactone display resistance. However, the cellular mechanisms driving cell death consequential to Sec61 inhibition remain poorly defined.

Mycolactone-sensitive proteins are still translated at the inhibited ribosome-translocon complex, but they are made in the cytosol rather being co-translationally translocated into the ER lumen. These mis-localised proteins are a potential source of proteotoxic stress, and we have shown that they are ubiquitinated and degraded by the proteasome, as well as by selective autophagy driven by an integrated stress response (ISR) rather than the classic ULK-dependent pathway (Hall *et al*, 2021). Inhibition of either autophagy or the ISR leads to rapid cell death (Ogbechi *et al*, 2018), suggesting that this is a protective response and that upregulation of autophagy flux has an important role in cell survival.

Our findings relating to autophagy induction, however, support a complex cellular phenotype that evolves over time. For instance, following 8hrs of exposure to mycolactone, there is clear indication of increased autophagic flux since increased puncta positive for early autophagic markers WIPI2, ATG16L1 and FIP200 are observed, as well as increased flux determined by assays with the tandem tag probe (mCherry-eGFP-LC3) (Pankiv *et al*, 2007). On the other hand, after 24hrs, evidence of autophagy inhibition starts to emerge, as cells begin to accumulate the adaptor protein SQSTM1/p62, which is normally consumed after autophagosome/lysosome fusion and builds up when late stage autophagy is inhibited (Klionsky *et al*, 2021). Even taking into account the increased production of SQSTM1/p62 by transcriptional and translational upregulation promoted by the ISR, this appears to represent a different phase of the response (Hall *et al*., 2021). We therefore postulated that the Sec61-dependent effects of mycolactone may impact lysosomal function over the longer term and that this is an important aspect of the mechanism of cell death.

## Results and Discussion

### Mycolactone disrupts lysosomal function

To assess the long-term impact of mycolactone on autophagic flux, we used the same tandem tag probe (mCherry-eGFP-LC3, Fig1A), but over an extended range of timepoints in HeLa cells (Fig1B). In cells with a functioning autophagy pathway, LC3-labelled autophagosomes form only transiently before fusing with the lysosome. Since the eGFP signal is quenched in lysosomes due to the reduced pH, these autolysosomes are typically identified by mCherry^+ve^ puncta (magenta in Fig1A&B), while autophagosomes yet to fuse with lysosomes are typically identified by mCherry^+ve^/eGFP^+ve^ puncta (white in Fig1A&B). Changes to flux can be observed when cells are exposed to saturating levels of the vATPase inhibitor BAFA1, inhibiting lysosomal acidification (Bowman *et al*, 1988; Klionsky *et al*., 2021; Pankiv *et al*., 2007).

Hence, as we previously observed, after 8hrs of exposure to mycolactone there were significantly increased numbers of mCherry^+ve^/eGFP^+ve^ (white) autophagosomes in the presence of BAFA1, confirming an early increase in autophagy flux. However, at later timepoints (24 and 32hrs) this effect was reversed. Notably, at these same later timepoints the number of mCherry^+ve^ lysosomes (magenta) also decreased, and by 32hrs was extremely low. At this stage, the relative abundance of mCherry^+ve^ and mCherry^+ve^/eGFP^+ve^ puncta expressed as a proportion (magenta:white) had also decreased significantly, with approximately 50% of puncta dual-labelled, suggesting an inhibition of fusion with lysosomes and/or an increase in lysosomal pH (Fig 1B, lower panels). However, since BAFA1 could still increase the number of mCherry^+ve^/eGFP^+ve^ (white) autophagosomes at 32hr, the effect was only partial. The tandem tag data suggests that after an initial increase in autophagic flux, long term exposure to mycolactone interferes with the later stages of the autophagy degradation pathway. This resolves the apparent discrepancy between our previous finding that mycolactone activates autophagy and an earlier report that mycolactone inhibits the pathway (Gama *et al*, 2014).

**Figure 1.**
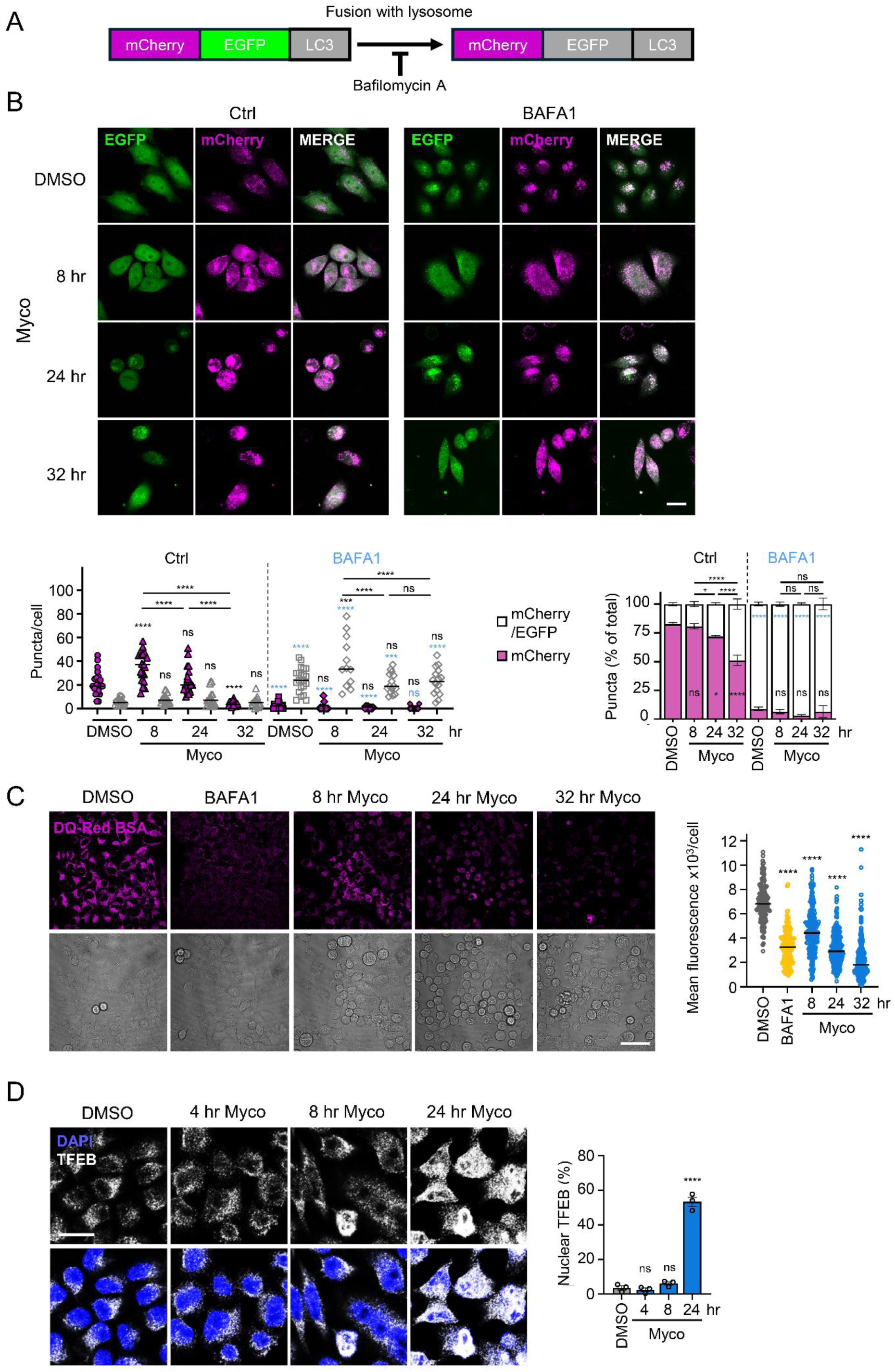
Mycolactone blocks late-stage autophagy, disrupts lysosomal function and activates TFEB signalling pathway. **A.** Schematic of LC3 tandem tag assay. **B-D.** HeLa cells were incubated with 0.05% DMSO for 32hr or 31.25ng/ml mycolactone (Myco) for various times as indicated with or without 300nM Bafilomycin A1 (BAFA1) for 4hr. **B.** Cells transiently transfected with pDEST-mcherry-eGFP-LC3B before treatment with mycolactone as described followed by live cell imaging, with the mCherry fluorescence pseudocoloured as magenta and eGFP as green. Quantitation shows the total number per cell (left hand graph, data representative of n=3 independent experiments; ordinary 2-way ANOVA) and relative proportions of magenta and white puncta (right hand graph, mean of n=3 biological replicates ± SEM; RM 2-way ANOVA) visualised in cells expressing mCherry-eGFP-LC3B. Scale bar =20µm. Data are compared to DMSO in the absence and presence of BAFA1 separately, and the effect of BAFA1 on each treatment is compared (indicated in blue). **C.** Cells were incubated with 20µg/ml DQ Red BSA for 4hr before washing and imaging for the unquenched signal. Scale bar =50µm. Quantitation of mean fluorescence per cell for at least 178 cells (data representative of n=3 independent experiments, ordinary 1-way ANOVA). **D.** Cells were fixed with 4% PFA, permeabilised, stained with anti-TFEB antibodies (grey) and counterstained with DAPI. Scale bar = 20µm. Graph shows mean % cells with nuclear TFEB staining from n=3 biological replicates ± SEM. ns; not significant, *; p<0.05, ***; p<0.001, ****; p<0.0001.

This effect was not due to cell death as cells remained ∼90% viable for up to 48hr in the presence of mycolactone, and significant levels of apoptotic death were only seen after 72hr (Fig S1A). Notably another structurally distinct Sec61 inhibitor, Coibamide A, induces a similar response, validating translocon inhibition as the cellular target driving the effect (Shi *et al*, 2021; Tranter *et al*, 2020).

One explanation for these observations could be loss of lysosomal function, since this is required to complete the process of autophagy. To determine whether mycolactone alters lysosomal proteolytic digestion capacity, HeLa cells were incubated with the DQ Red BSA, a fluorogenic protease substrate which is de-quenched following proteolysis, yielding a strong fluorescent signal in lysosomes. In control cells, high levels of fluorescence were detected in all cells (Fig 1C). As expected, this signal was blocked by incubation with BAFA1 due to loss of lysosome acidification. Mycolactone caused a progressive decrease in DQ red BSA fluorescence which was statistically significant as early as 8hr after exposure and even lower than that seen in BAFA1-treated cells by 32hrs. This is not specific to HeLa cells as similar results were obtained in primary murine bone marrow derived macrophages as well as human macrophage-like cells (PMA-activated THP-1 cells) (Figure S1C,D). This shows that Sec61 inhibition by mycolactone causes a general suppression of lysosomal function.

The transcription factor TFEB controls autophagy and lysosomal biogenesis by upregulating genes in the Coordinated Lysosomal Expression and Regulation (CLEAR) network (Sardiello *et al*, 2009). TFEB activation and nuclear translocation follows loss of mTORC-dependent phosphorylation that can be triggered by multiple pathways that impact autophagy, such as nutrient deprivation and metabolic or oxidative stress, but can also be activated in response to lysosomal dysfunction caused by lysosomal storage disorders, membrane damage, calcium loss or changes in luminal pH and may be used as an indicator of lysosomal stress (Pena-Llopis *et al*, 2011; Raben & Puertollano, 2016). In control cells, TFEB was predominantly cytoplasmic and little change could be seen up to 8hr after mycolactone addition but by 24hr over 50% of nuclei were positive for TFEB (Fig 1D). This confirms our earlier findings that mTOR inactivation is a relatively late-stage event in response to mycolactone and suggests that the loss of function following long-term exposure to mycolactone causes lysosomal stress (Hall *et al*., 2021; Ogbechi *et al*., 2018).

### Mycolactone rapidly downregulates the V-ATPase accessory subunits ATP6AP1 and ATP6AP2

Lysosome function depends heavily on luminal and membrane proteins that are dependent on the Sec61 for their translocation into the ER. Such constitutively expressed proteins have been shown to be depleted at their turnover rate following Sec61 inhibition (Ogbechi *et al*, 2015). However, the rate of depletion of individual proteins is difficult to predict and can change under stress conditions. In our recent proteomic study of human dermal microvascular endothelial cells (HDMEC) (Hsieh *et al*., 2025), we detected 138 lysosome-associated proteins, of which 24 showed reduced expression after 24hrs mycolactone exposure (PRIDE Project PXD037489). However, since only two proteases, legumain and Cathepsin S, were significantly downregulated at this point, loss of proteases alone seems to be insufficient to explain the impact of mycolactone on lysosome function.

We therefore turned our attention to the V-ATPase, which is responsible for the acidic pH of lysosomes (as well as the Golgi and endosomal system) and thus regulates the activity of many lysosomal enzymes. The V-ATPase is a complex made up of two multi-subunit domains, the cytosolic V_1_, responsible for ATP hydrolysis, and the membrane-integral V_0_, which translocates protons across the membrane (Fig 2A) (Abbas *et al*, 2020; Nelson & Harvey, 1999; Wang *et al*, 2020). The proteins making up the V-ATPase complex, and their predicted and actual sensitivity to mycolactone are shown in Fig 2B. The cytosolic V_1_ proteins and the multipass components of the V_0_ complex are not predicted to be targets of mycolactone due to their topology, and their abundance was not significantly changed after 24hrs mycolactone exposure in HDMEC (Fig 2B). Indeed, this was confirmed for ATP6V0D1 and ATP6V1H, representing each category in HDMEC and HeLa (Fig 2C).

**Figure 2.**
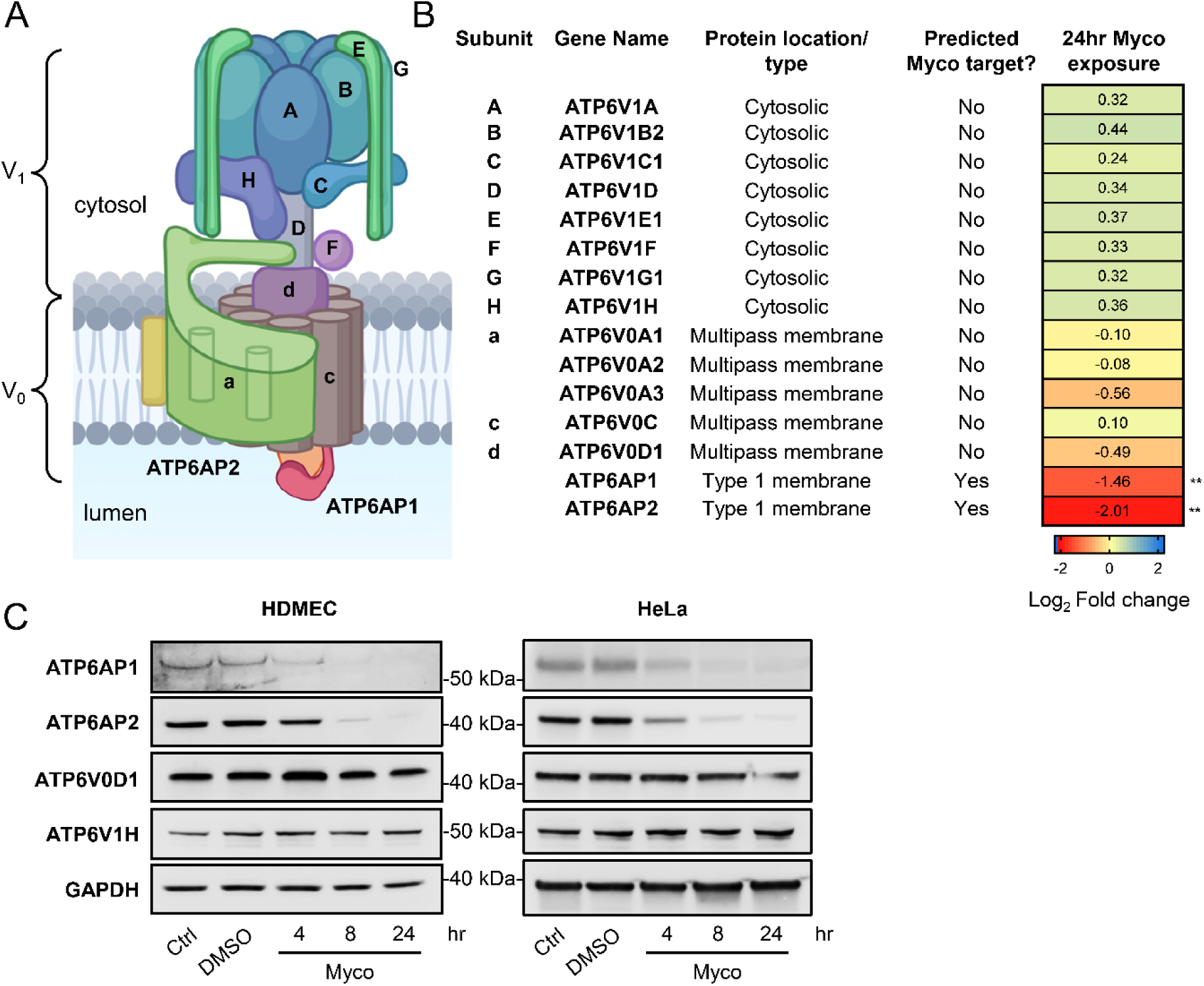
Mycolactone targets the ATP6A1 and ATP6A2 subunits of the Vacuolar ATPase. **A.** Cartoon structure of V-ATPase complex, depicting the cytosolic V_1_ complex responsible for ATP hydrolysis and the membrane embedded V_0_ which pumps protons across the membrane. Image generated using Biorender. **B.** V-ATPase subunits detected in our HDMEC membrane proteome (PRIDE Project PXD037489)(Hsieh *et al*., 2025) showing predicted sensitivity to mycolactone based on location and topology combined with the known specificity of mycolactone towards different Sec61 substrates, along with a heat map of Log2 fold-change in V-ATPase subunit expression following mycolactone exposure for 24hr. **; p<0.01. **C.** Immunoblots from HDMEC incubated with 0.02% DMSO or 10ng/ml mycolactone and HeLa cells incubated with 0.05% DMSO for 24hr or 31.25ng/ml mycolactone for various times and stained with the indicated antibodies. GAPDH is included as a loading control. Approximate molecular weights in kDa are indicated for each target. Images are representative of at least n=2 independent experiments.

However, the two single-pass type I membrane proteins ATP6AP1 and ATP6AP2 are predicted targets of mycolactone and are both significantly depleted following 24hrs exposure (Fig 2B). We explored the kinetics of this depletion and found that both ATP6AP1 and ATP6AP2 are rapidly lost in the presence of mycolactone in primary HDMEC, HeLa (Fig2C) and PMA-differentiated THP-1 (Fig S2) cells, with a reduction in each detectable as early as 4hr post-exposure.

ATP6AP1 and ATP6AP2 are accessory proteins of the V-ATPase. While they are not directly involved in proton translocation, they are essential for biogenesis of functional V-ATPase complexes and are important for both lysosome function and production (Abbas *et al*., 2020; Nelson & Harvey, 1999; Wang *et al*., 2020). The two appear to act in concert, with ATP6AP1 acting as a hub around which the V_0_ membrane components assemble in the ER, controlling both stability and trafficking of the complex (Abbas *et al*., 2020; Guida *et al*, 2018; Wang *et al*., 2020). Their knockdown, deletion or mutation has been shown to reduce V_0_ protein levels and increase lysosomal and endosomal pH (Pareja *et al*, 2018). Moreover, depletion of either ATP6AP1 or ATP6AP2 alone can reduce the number of lysosomes, decrease cargo degradation and disrupt autophagy (Pareja *et al*., 2018). ATP6AP1 may even play a direct role in autophagosome-lysosome fusion via interaction with Rab7 and the HOPS complex (Yan *et al*, 2025). Genetic disorders caused by mutations in ATP6AP1 and ATP6AP2 affect protein glycosylation and have pleiotropic effects on multiple tissues while altered expression and loss of function mutations are seen in many cancers (Jansen *et al*, 2016; Pareja *et al*., 2018; Qi *et al*, 2022; Rujano *et al*, 2017). Their downregulation by mycolactone would thus be expected to affect multiple cellular functions but have a particularly heavy impact on lysosomes and lysosome dependent processes.

### Mycolactone reduces association of V1 complex with lysosomes

SidK is a bacterial toxin which binds specifically to the cytoplasmic ATP6V1A subunit of the V-ATPase (Xu *et al*, 2010) and so expression of fluorescently-tagged truncation mutants of SidK can be used as probes to localise and quantify functional V-ATPase complexes in cells (Maxson *et al*, 2022). In order to investigate the effect of mycolactone on the whole V-ATPase complex, we used a SidK-mCherry construct (magenta in Fig 3A) and examined co-localisation with LAMP1 (green in Fig3A). In control cells, some regions were positive for SidK-mCherry but not LAMP1, likely V-ATPase associated with Golgi and endosomal membranes. However, the majority of LAMP1^+ve^ compartments coincided with SidK-mCherry (white in Fig4A). The same pattern of staining was observed in cells exposed to mycolactone for 8hr and 24hr. However, at 32hr mycolactone exposure, there was little overlap of signals and cells exhibited a redistribution of both LAMP1 and SidK-mCherry positivity from the perinuclear region to the cell periphery. The effect of mycolactone on SidK localisation was not just a consequence of altered LAMP1 expression as a similar effect was observed in cells labelled with Alexa 647 Dextran, which accumulates in lysosomes and provides and independent measure of lysosome numbers (Fig S4A).

**Figure 3.**
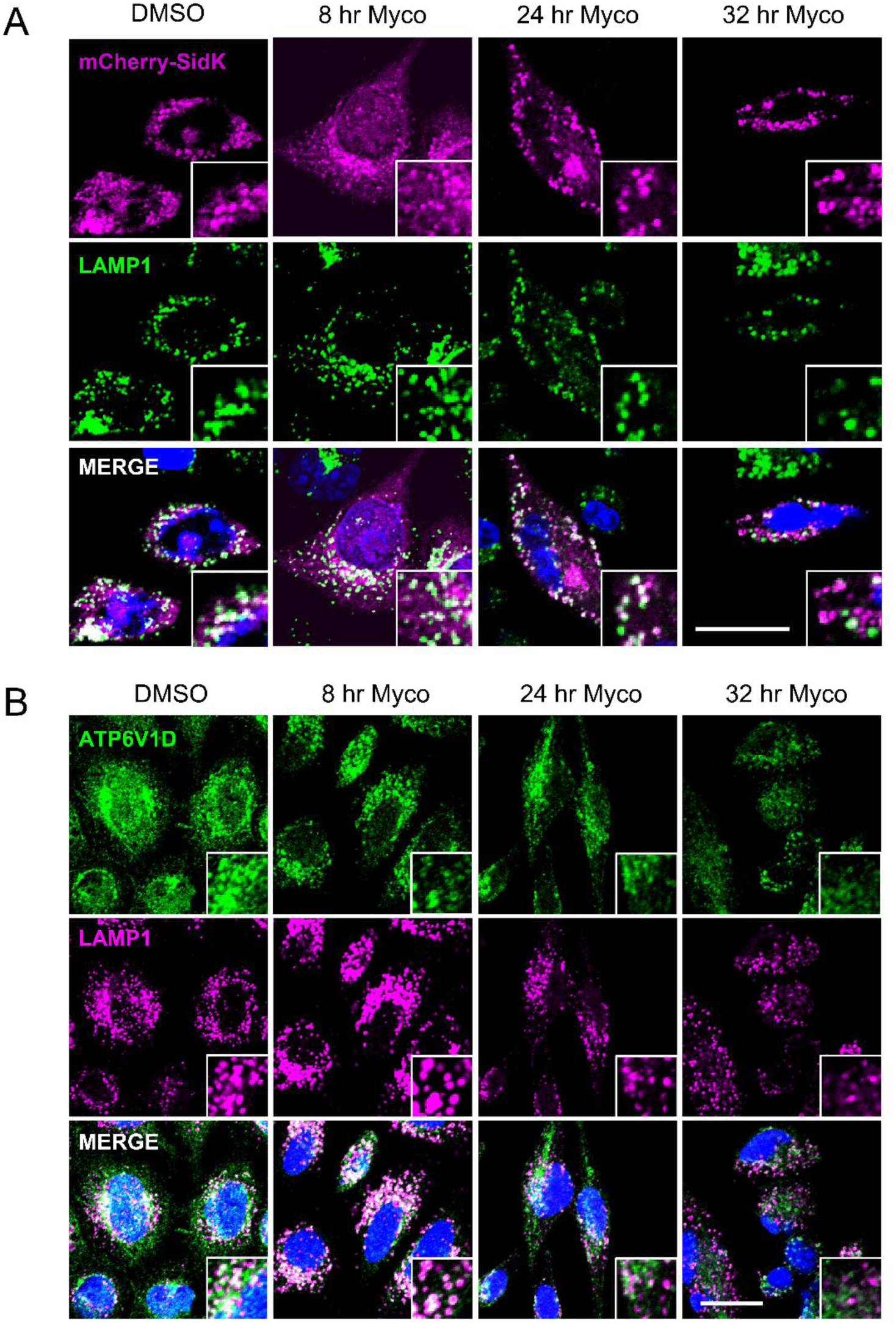
Reduced association of V-ATPase V_1_ with lysosomes. HeLa cells were incubated with 0.05% DMSO for 32hr or 31.25ng/ml mycolactone (Myco) for various times. All cells were counterstaining with DAPI; scale bar = 20µm. Images representative of n=3 independent experiments. **A.** Cells were transiently transfected with pYMA1_SidK_mCherry (pseudocoloured in magenta) prior to the experiment, and fixed/stained with anti-human LAMP1 antibodies (green). **B.** Cells were fixed/stained with anti-ATP6V1D (green) and anti-human LAMP1 antibodies (magenta).

To rule out any potential confounding effects from over-expression of SidK-mCherry, which can itself have an impact on lysosomal function (Maxson *et al*., 2022), we examined the co-localisation of LAMP1 with ATP6V1D (to detect membrane associated V_1_, Fig 4B). In control cells, most LAMP1 co-localised with ATP6V1D as expected. This did not change after 8hrs of mycolactone exposure but, at 24hr, both the level of membrane associated ATP6V1D and the proportion colocalising with LAMP1 were reduced and by 32hr there was little overlap between the signals, confirming that the V_1_ complex no longer associated with lysosomes.

**Figure 4.**
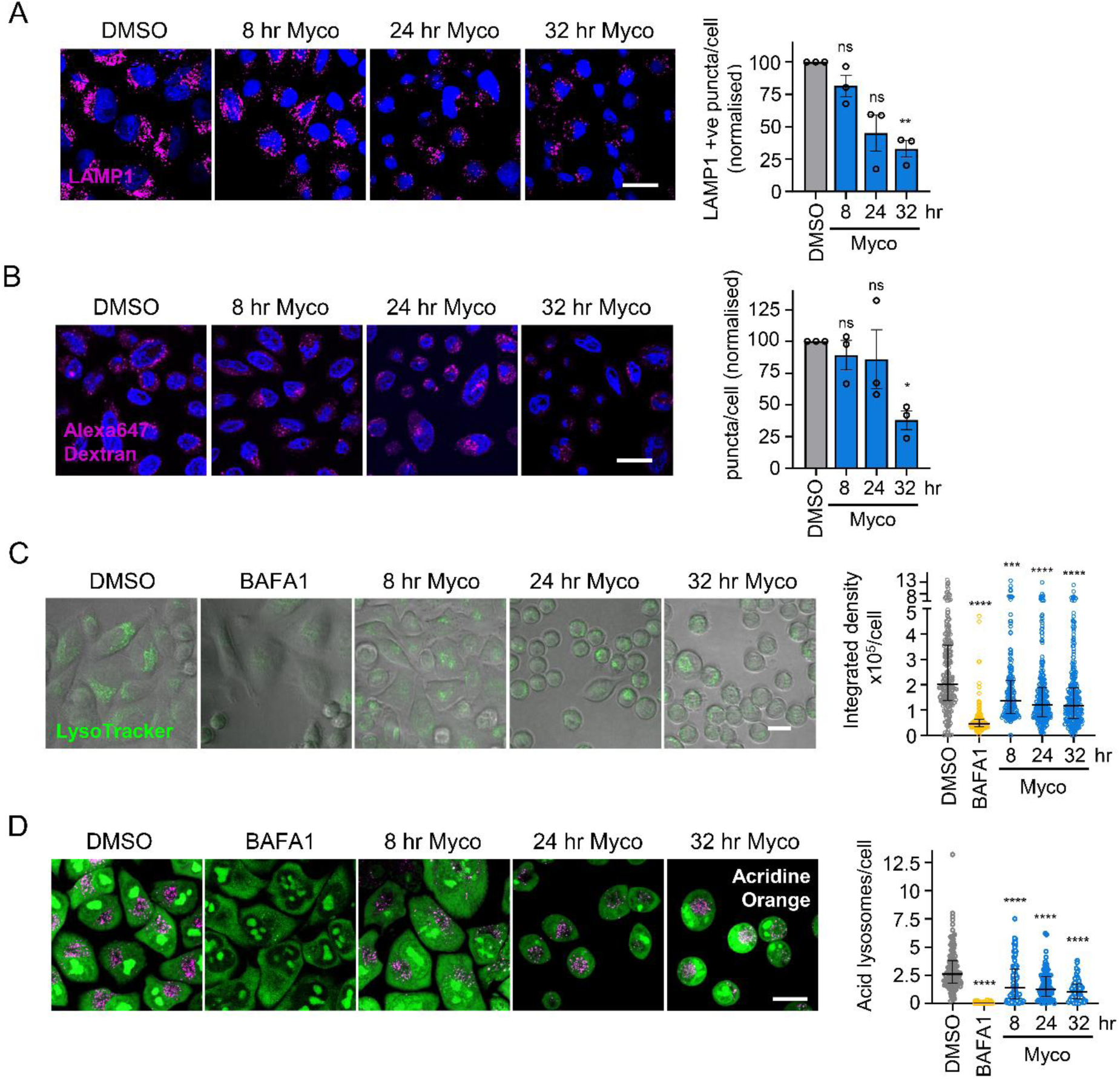
Prolonged exposure to mycolactone leads to a reduction in lysosome number and acidity. **A.** HeLa cells were incubated with mycolactone, fixed with 4% PFA, permeabilised, stained with anti-LAMP1 antibodies (magenta) and counterstained with DAPI. Quantitation of mean LAMP1 positive puncta per cell. Data represents the normalised mean of 3 independent experiments ± SEM, one-sample t-test. **B.** HeLa cells pulsed with 0.1mg/ml Alexa 647 dextran for 3hr and chased for 1hr before fixing and counterstaining with DAPI. Quantitation of mean Alexa 647 positive puncta per cell. Data represents the normalised mean of n=3 independent experiments ± SEM (one-sample t-test). **C.** Hela cells incubated as described above, then incubated with 75nM Lysotracker Green DND-26 for 10min before washing. Cells were re-incubated 10min then imaged. Quantitation of integrated density of Lysotracker fluorescence per cell for at least 330 cells. Data is representative of n=3 independent experiments showing median ± IQR (Kruskal-Wallis test). **D.** HeLa cells were incubated as described then incubated with 2µM acridine orange for 30min before washing and imaging. Acid lysosomes are pseudocoloured magenta, whereas the green is observed where the acridine orange remains in non-acidic compartments. Quantitation of acid lysosomes for at least 61 cells is shown. Data is representative of n=3 independent experiments showing median ± IQR, Kruskal-Wallis test. All scale bars = 20µm. ns; not significant, *; p<0.05, **; p<0.01, ***; p<0.001, ****; p<0.0001.

At 32 hours, SidK-mCherry remained associated with vesicular compartments, and we hypothesised that this could be the Golgi, since this is the location of V-ATPase assembly. However, we also observed a loss over time of co-localisation between SidK-mCherry, as well as ATP6V1D, with the Golgi marker Giantin, becoming pronounced at 32hr (Fig S4B and Fig S4C respectively). Therefore, it is not clear at the current time what these compartments are. Nevertheless, these results suggest that levels of functional V-ATPase gradually decline over time following exposure to mycolactone. The difference in kinetics between loss of ATP6AP1 and ATP6AP2 expression and the V_0_ complex suggest the accessory proteins may be made in excess and it is not until they are completely lost, and pre-existing complexes are also turned over, that the full impact on V-ATPase complex formation can be seen.

### Mycolactone causes a decrease in lysosome number and acidity

The effects we have observed in assays with the LC3 tandem tag, DQ BSA and V_1_ localisation could reflect a reduction in lysosomal number, impaired acidification or both. In order to tease these apart, we first enumerated lysosomes and found mycolactone caused a significant decrease in LAMP1^+ve^ vesicles in HeLa cells that became significant after 32hr (p<0.01; Fig 4A). Since LAMP1 is a signal peptide-containing protein, and therefore a predicted direct target of mycolactone, we also assessed lysosome number by labelling with the dye Alexa647-dextran (Fig 4B). Here, a significant reduction in Alexa647-dextran^+ve^ puncta (p<0.05) was also seen at 32hrs, confirming that total lysosome numbers are indeed reduced by prolonged exposure to mycolactone.

It is possible the reduced levels of LAMP1 could, on its own, explain the fall in the lysosome numbers, as LAMP1 has been implicated in lysosome biogenesis and has been shown to regulate pH via interaction with the cation channel TMEM175 (Zhang *et al*, 2023). However, in contrast to our *in vitro* findings, lysosomal proteolytic activity in mice deficient in LAMP1 has been reported to be unchanged (Andrejewski *et al*, 1999) and while this offers an explanation for the inhibition of late-stage autophagy at 32hr, it does not explain the reduced DQ Red BSA proteolysis at earlier timepoints (8hrs, Fig 1C). Irrespective of the underlying mechanism, the results show that Sec61 inhibition affects lysosome biogenesis/turnover.

Depletion of ATP6AP1 and/or ATP6AP2 leads to defects in lysosomal acidification (Guida *et al*., 2018; Pareja *et al*., 2018; Rujano *et al*., 2017; Yan *et al*., 2025). To determine whether mycolactone has an impact on lysosomal pH we labelled cells with LysoTracker Green DND-26, which accumulates in acidic compartments. In control cells, LysoTracker staining could be seen in most cells, predominantly in the perinuclear region, while the signal was profoundly depressed on inclusion of BAFA1 to block V-ATPase activity, as expected (Fig 4C). At 8 hours, LysoTracker could still be detected in many mycolactone-exposed cells but the median was reduced compared to the control (p<0.001). After 24hr, while some cells still exhibited strong fluorescence, others showed very low LysoTracker positivity. The proportion of these low stained cells increased at 32hr but did not reach the level of inhibition seen with BAFA1.

We hypothesised that the results with LysoTracker were due to the effect of mycolactone on lysosomal pH. To test this, we incubated cells with acridine orange, a cell permeable dye which accumulates in lysosomes. Acridine orange emits green fluorescence in its monomeric form but, in an acidic environment dimerises, causing a shift to red fluorescence emission, as seen in control conditions (Fig 4D). This was blocked by BAFA1, as expected. As with LysoTracker, on exposure to mycolactone, the number of acidic lysosomes per cell decreased over time (p<0.001). Taken together, the results show that mycolactone reduces the number of acid lysosomes in cells and, since this precedes the reduction in total lysosomes, this may be due at least in part to impaired acidification following loss of ATP6AP1 and ATP6AP2.

*M. ulcerans* is a mycobacterium that infects macrophages as part of its lifecycle. Since the endolysomal system is critical for the host defence against mycobacteria, we investigated whether mycolactone also reduced LysoTracker accumulation in BMDM and THP-1 cells (Fig S4). This was indeed the case, suggesting that the ability to down-regulate the V-ATPase may play a role in the infection cycle of *M. ulcerans* in Buruli ulcer disease. This provides an explanation for previous reports that showed reduced accumulation of LysoTracker dye in the phagosomes of macrophages infected with mycolactone-positive *M. ulcerans* (Torrado *et al*, 2010). We propose that this explains how *M. ulcerans* can grow in host macrophages despite the well-described genome reduction of this pathogen. Deletions and loss of function in the locus encoding ESX1, a secretion system required for lysosome escape that is essential for virulence of *M. tuberculosis* and *M. marinum* (Stinear *et al*, 2007; Torrado *et al*, 2007), mean that *M. ulcerans* must use alternative strategies for intra-macrophage survival.

Hence, our findings demonstrate a novel virulence mechanism that would have a profound impact on host-pathogen interactions in Buruli ulcer. Our working model involves dysfunction of the endolysosomal system following reduced generation of acidic lysosomes due to the inability to form functional V-ATPase complexes, with the loss of ATP6AP1 and ATP6AP2 as the mechanism driving these changes. The delayed onset of loss of acidic lysosomes, despite the rapid loss of ATP6AP1 and ATP6AP2, is likely due to the rate of turnover of pre-existing V-ATPase complexes. Although there are other key lysosomal proteins downregulated by mycolactone, the loss of these two proteins alone is sufficient to cause the phenotype we observe, due to their critical role in V-ATPase complex assembly.

Interestingly, ATP6AP1 has recently also been shown to be a guanine nucleotide exchange factor for Rheb, a component of mTORC1 (Feng *et al*, 2024). Thus, its depletion may be a factor in the nuclear translocation of TFEB seen following mycolactone exposure. However, the consequent activation of lysosomal stress pathways is ineffective, since the cells are no longer able to respond by upregulating lysosome biogenesis. We propose that this, combined with the block in late-stage autophagy, further compromises the capacity of the cell to cope with the proteostatic stress caused by Sec61 inhibition. Therefore, the virulence mechanism not only has a direct impact on bacterial survival in the macrophage, it would also contribute to the death of cells surrounding the infection foci and the consequent tissue necrosis seen in Buruli ulcer.

The effects of Sec61 inhibition on lysosomes has broader implications, since Sec61 inhibitors are under investigation as treatments for cancer, viral infections and rheumatoid arthritis (Domenger *et al*, 2022; Heaton *et al*, 2016; Rehan *et al*, 2023). The long-term impact of mycolactone on lysosomal function may be a key component of its cytotoxicity that can be exploited, but could also represent a source of unwanted side effects in Sec61 targeting therapies.

## Methods

### Mycolactone

For all experiments, we used synthetic mycolactone A/B, which was generously donated by Prof. Yoshito Kishi (Harvard University). To control for potential impact of the DMSO solvent on cell function, DMSO diluted equivalently was used; typically this was 0.02%.

### Cell culture and treatment

HeLa and L929 cells were maintained in DMEM supplemented with 10% FBS at 37°C and 5% CO2. To generate conditioned medium, L929 cells were cultured to confluency, then maintained for 3 further days. Supernatants were collected and spun at 300 x g for 5min, then filtered and stored at -70⁰C. Juvenile, single donor human microvascular endothelial cells (HDMEC) were cultured in hVEGF containing Endothelial cell growth medium 2 (Promocell) at 37°C and 5% CO_2_. Cells were routinely seeded at a concentration of 1 x 10^4^/cm^2^ in 25cm^2^ or 75cm^2^ flasks for no more than 15 population doublings. THP-1 cells were cultured in suspension in RPMI supplemented with 10% FBS at 37°C and 5% CO_2_. Cell density was maintained between 0.1 and 10^5^ cells/ml. For differentiation, cells were plated out at a density of 5 x 10^5^/ml in the presence of 20ng/ml PMA for 72hr. Medium was replaced and the cells allowed to rest for 24hr before experiments. BMDM derived from C57BL/6J mice were differentiated for 7 days in 80% RPMI supplemented with 10% FBS/20% L929 conditioned medium. After harvesting cells were plated out at a density of 5 x 10^5^/ml. Mycolactone was used at a concentration of 31.25ng/ml for HeLa cells, THP-1 and BMDM and at 10ng/ml for HDMEC. Bafilomycin A1 was used at 300nM for HeLa cells and 200nM for all other lines.

### Transfections

The tandem tag vector pDEST-mCherry-eGFP-LC3B was a gift from Terje Johansen and pYMA1_SidK_mCH was a gift from John Rubinstein (Addgene plasmid # 184906; http://n2t.net/addgene:184906; RRID:Addgene_184906). Cells in 80% confluent 6-well plates were transfected overnight with 2µg plasmid DNA then trypsinised and plated onto coverslips for co-staining with antibodies or IBIDI chamber slides for live cell imaging. Cells were incubated overnight then with DMSO, mycolactone or BAFA1 as described then pDEST-mCherry-eGFP-LC3B transfected cells were imaged live while pYMA1_SidK_mCH transfected cells were fixed for immunofluorescent staining.

### Immunoblotting

Immunoblotting was carried out according to standard methods as described in Ogbechi *et al*. (Ogbechi *et al*., 2018). Cells were plated onto 6 well plates, incubated with mycolactone for various times then lysed directly in 1 x Laemmli SDS sample buffer (with sonication to degrade genomic DNA). Proteins were separated on 4-20% pre-cast gels (BioRad) and blotted onto Immobilon PVDF membranes (Merck). Blots were blocked with 5% milk powder in TBS/0.05% Tween 20. Primary antibodies were diluted in 3% BSA/TBS/0.05% Tween 20). Antibodies used in this study were: Anti-ATP6AP1 (Santa Cruz Biotechnology, sc-81886); anti-ATP6AP2 (Bio-Techne, NBP1-90820); anti-ATP6V0D1 (Proteintech, Proteintech: 18274-1-AP); anti-ATP6V1H (Cambridge Bioscience, A304-308A) and anti-GAPDH (Fisher Scientific, 10515025). Secondary antibodies were anti-rabbit-HRP (GE Healthcare, NA934V) and anti-mouse-HRP (GE Healthcare, NA931V) and protein bands were visualised using Immobilon Western Chemiluminscence Substrate (Fisher Scientific). Images were developed on a Fusion FX Imager (Vilber-Lourmatclean), which provides a warning if the areas of the image are saturated.

### Immunofluorescence

Immunofluorescent imaging was carried out according to standard methods as described (Hall *et al*., 2021). Cells were plated onto sterile glass coverslips and incubated overnight before experiments. Cells were either fixed with ice-cold methanol and blocked with 2% BSA/PBS overnight before staining or fixed with 4% paraformaldehyde in 200mM Hepes pH7.5 then were permeabilised wit 0.25% Nonidet P-40 alternative in NETGEL buffer (150 mM NaCl, 5mM EDTA, 50 mM Tris-Cl, pH 7.4, 0.05% Nonidet P-40 alternative, 0.25% gelatin and 0.02% sodium azide) and blocked with NETGEL buffer. Antibodies used in this study were: anti-TFEB (Cambridge Bioscience, A303-673A); anti-human LAMP1 (Bio-Techne, MAB4800): anti-ATP6V1D (Abcam, ab157458) and anti-Giantin (Abcam, ab80864). Secondary antibodies were: Alexa Fluor 594 goat anti-rabbit (A11012); Alexa Fluor 488 donkey anti-mouse (A21202); Alexa Fluor 488 goat anti-rabbit (A11034); Alexa Fluor 647 goat anti-rabbit (A21244) and Alexa Fluor 647 goat anti-mouse 647 (A21235), all from Fisher Scientific. Slides were counterstained with 0.1μg/ml DAPI and mounted with Polymount (Tebu-Bio).

### DQ Red BSA assay

Cells were plated onto IBIDI 8-well chamber slides and incubated overnight then incubated with 0.05% DMSO for 32hr, 300nM Bafilomycin A1 (BAFA1) for 4hr or 31.25ng/ml mycolactone (Myco) for various times. DQ Red BSA (20μg/ml) (Fisher Scientific) was added in Optimem (Fisher Scientific) 4hr before the end of the experiment. Cells were gently washed three times with Live Cell Imaging Medium (Fisher Scientific) and returned to the incubator for 5min before imaging.

### Alexa Fluor 647 Dextran labelling

Cells plated onto coverslips were incubated overnight then with 0.05% DMSO for 32hr, 300nM Bafilomycin A1 (BAFA1) for 4hr or 31.25ng/ml mycolactone (Myco) for various times. Alexa Fluor 647 Dextran 10,000 (0.1mg/ml) was added for three hours, then cells were washed and incubated for 1hr in fresh medium. Cells were fixed and mounted on Polymount for imaging

### Lysotracker labelling

Cells were plated onto IBIDI 8-well chamber slides and incubated overnight then incubated with 0.05% DMSO for 32hr, 300nM Bafilomycin A1 (BAFA1) for 4hr or 31.25ng/ml mycolactone (Myco) for various times then incubated for 10min with Lysotracker Green DND-26 (Fisher Scientific). Cells were gently washed three times with Live Cell Imaging Medium (Fisher Scientific) and returned to the incubator for 10min before imaging live.

### Acridine orange

Cells were plated onto IBIDI 8-well chamber slides and incubated overnight then incubated with 0.05% DMSO for 32hr, 300nM Bafilomycin A1 (BAFA1) for 4hr or 31.25ng/ml mycolactone (Myco) for various times. Acridine orange (2μM, Bio-Rad) was added for 30min, cells were washed three times with Live Cell Imaging Medium (Fisher Scientific) and imaged live.

### Imaging and analysis

All imaging was carried out on a Nikon Eclipse TI confocal microscope. All image analysis was carried out using ImageJ software. Puncta were counted manually for the tandem tag analysis and using the Analyse Particles function for all other assays. The Measure function was used to quantify mean and integrated density fluorescence per cell.

### Statistical analysis

All data were analysed using GraphPad Prism Version 10.4.1. Data were analysed using a one-sample t-test, or one- or two-way ANOVA using pairing if appropriate. Where data were not normally distributed, a non-parametric test (Kruskal-Wallis) was used. An appropriate correction for multiple comparisons (either Dunnett’s, Tukey’s or Dunn’s) was performed if possible.

## Data Availability

This study includes no data deposited in external repositories.

## Acknowledgements

We are extremely grateful to Prof Yoshito Kishi (Harvard University, USA) for the gift of synthetic mycolactone A/B and Dr Terje Johansen (University of Tromso, Norway) for the Tandem Tag pDEST-mCherry-eGFP-LC3B plasmid. pYMA1_SidK_mCH was a gift from John Rubinstein (Addgene plasmid # 184906). This work is funded by MRC project grant MR/W02618X/1. KO-B is supported by a University of Surrey Vice-Chancellor’s Award.

## Author Contributions

Conceptualization, BH and RS; methodology, BH; investigation, BH and KO-B; funding acquisition, BH and RS; project administration, RS; supervision, BH and RS; writing, BH and RS.

## Supplementary Figure Legends

**Figure S1.**
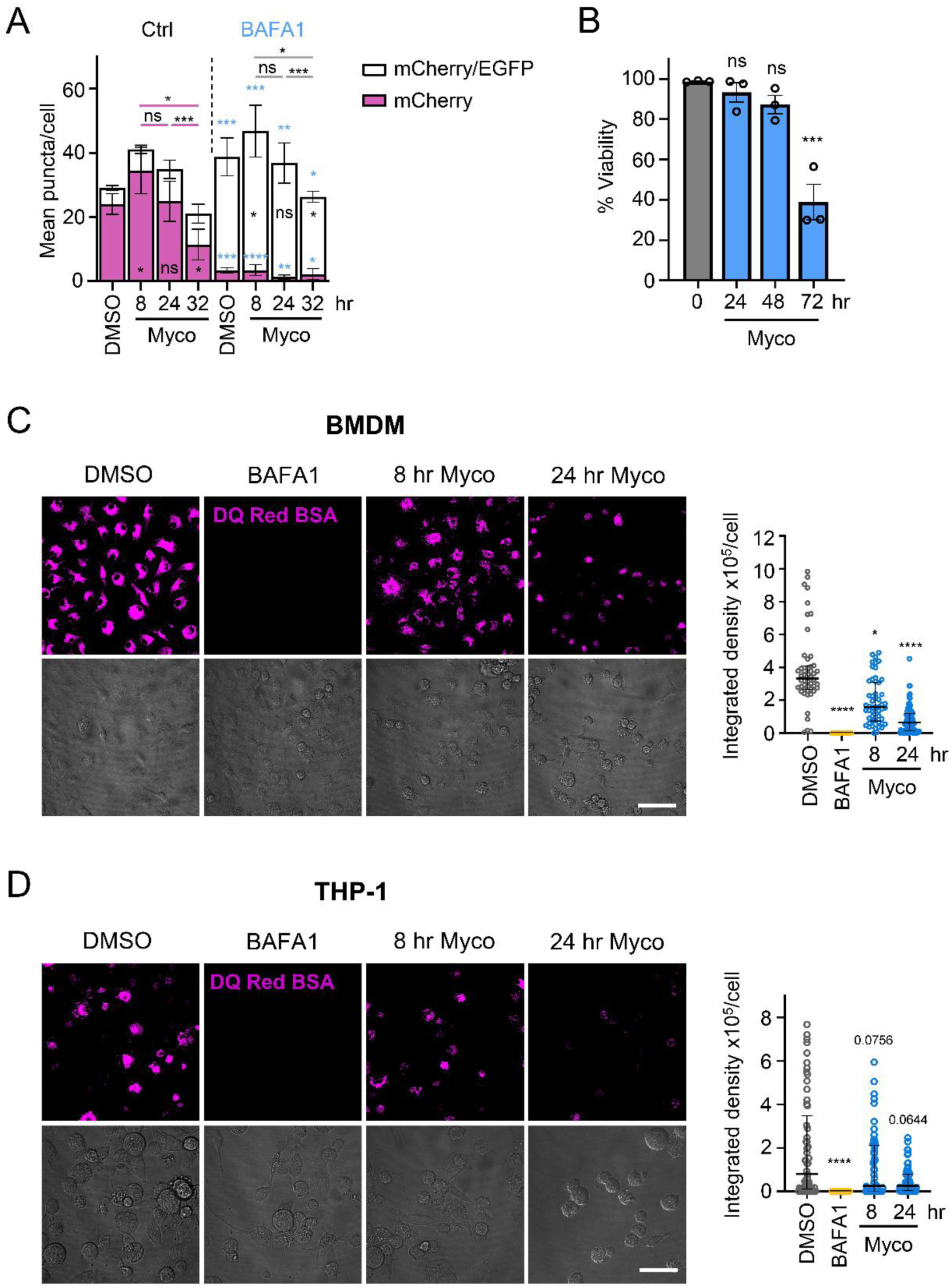
Mycolactone blocks late-stage autophagy, disrupts lysosomal function and activates TFEB signalling pathway. **A.** Cells were transiently transfected with pDEST-mcherry-eGFP-LC3B before treatment with mycolactone as described in Fig1A. Quantitation shows the total number of magenta and white puncta per cell. Data represents the mean of 3 independent experiments ± SEM analysed by RM 2-way ANOVA. Data are compared to DMSO in the absence and presence of BAFA1 separately, and the effect of BAFA1 on each treatment is compared (in blue). **B.** HeLa cells were incubated with 0.05% DMSO or 31.25ng/ml mycolactone (Myco) for various times, stained with CellEvent and propidium iodide and imaged. Data represents the % viable cells at each time point and is the mean of n=3 independent experiments ± SEM (RM one-way ANOVA). **C.** BMDM were seeded onto IBIDI 8-well chamber-slides and incubated with 0.05% DMSO for 24hr, 31.25ng/ml mycolactone (Myco) for various times or 200nM Bafilomycin A1 (BAFA1) for 4hr. DS Red BSA was added for 4hr before washing and imaging. Scale bar =50µm. Data represents median DS Red BSA fluorescence per cell for at least 48 cells ± IQR. Data representative of n=3 independent experiments (Kruskal-Wallis test). **D.** THP-1 cells were plated onto IBIDI 8-well chamber slides in the presence of 20ng/ml PMA. After 72hr medium was replaced and cells incubated for 24hr before adding 0.05% DMSO for 24hr, 31.25ng/ml mycolactone (Myco) for various times or 200nM Bafilomycin A1 (BAFA1) for 4hr. DS Red BSA was added for 4hr before washing and imaging. Scale bar =50µm. Data represents median DS Red BSA fluorescence per cell for at least 53 cells ± IQR. Data representative of n=3 independent experiments (Kruskal-Wallis test). ns; not significant, *; p<0.05, **; p<0.01, ***; p<0.001, ****; p<0.0001 or as indicated.

**Figure S2.**
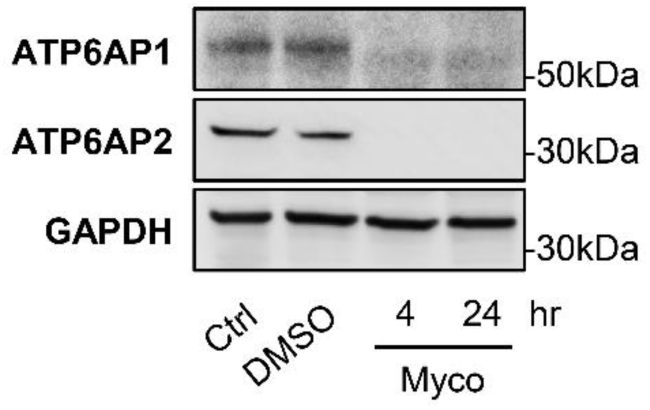
Mycolactone targets the ATP6A1 and ATP6A2 subunits of the Vacuolar ATPase. Immunoblot of THP-1 cell lysates incubated with 0.05% DMSO or mycolactone for various times and stained with anti-ATP6AP1 and -ATP6AP2 with anti-GAPDH is used as a loading control. Data is representative of n=3 independent experiments.

**Figure S3.**
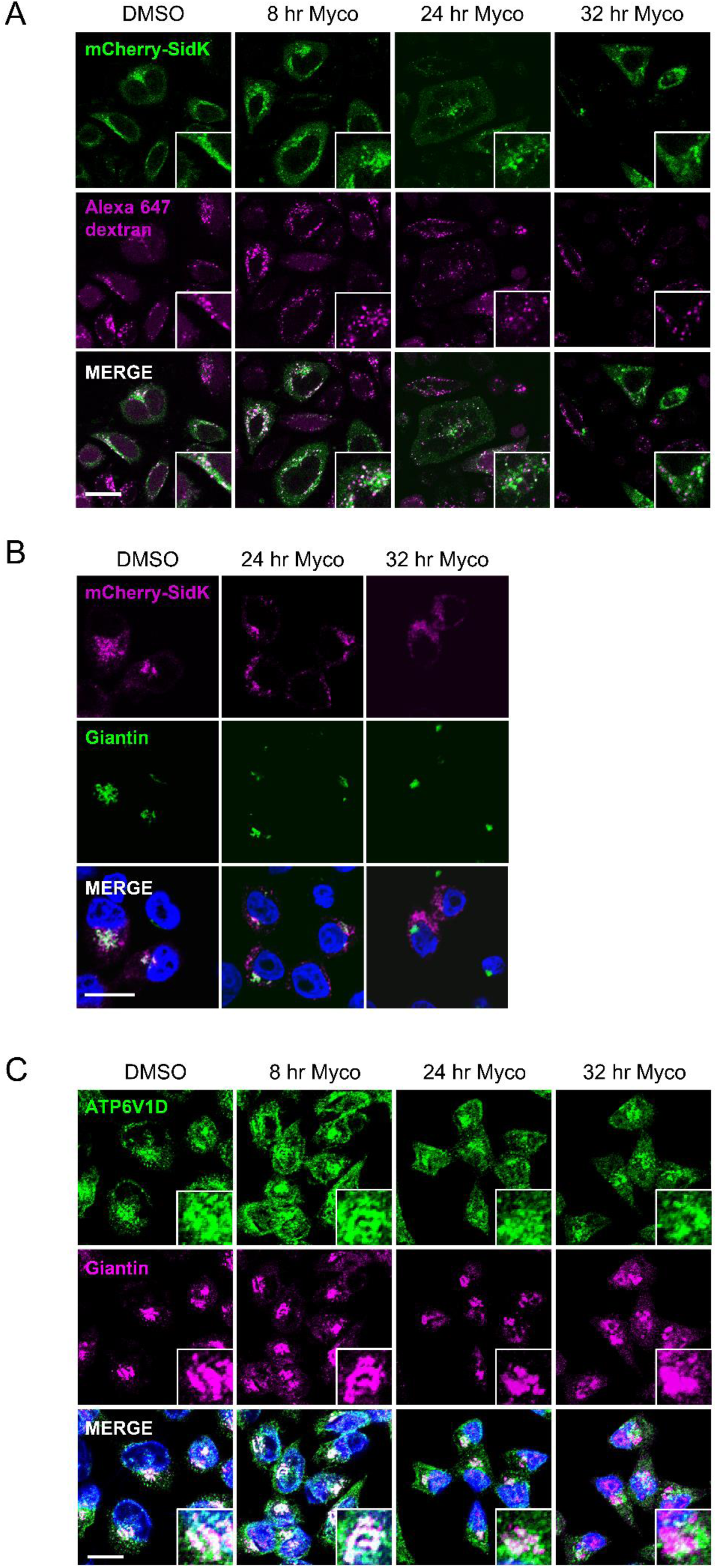
Reduced association of V-ATPase V_1_ with lysosomes. **A.** HeLa cells transiently transfected with pYMA1_SidK_mCH were incubated with 0.05% DMSO for 32hr or 31.25ng/ml mycolactone (Myco) for various times. Cells were pulsed with 0.1mg/ml Alexa 647 dextran (magenta) for 3hr then chased for 1hr before fixing. mCherry Sidk is pseudocoloured in green. Data is representative of n=2 independent experiments. **B.** HeLa cells transiently transfected with pYMA1_SidK_mCH were incubated with 0.05% DMSO for 32hr or 31.25ng/ml mycolactone (Myco) for various times before fixing and staining with anti-human Giantin antibodies (green) and counterstaining with DAPI with mCherry Sidk pseudocoloured in magenta. Scale bar = 20µm. Images are representative of n=3 independent experiments. **C.** HeLa cells were incubated with 0.05% DMSO for 32hr or 31.25ng/ml mycolactone (Myco) for various times, fixed with methanol, blocked and stained with anti-ATP6V1D (green) and anti-Giantin antibodies (magenta) then counterstained with DAPI. Scale bar = 20µm. Images are representative of n=2 independent experiments.

**Figure S4.**
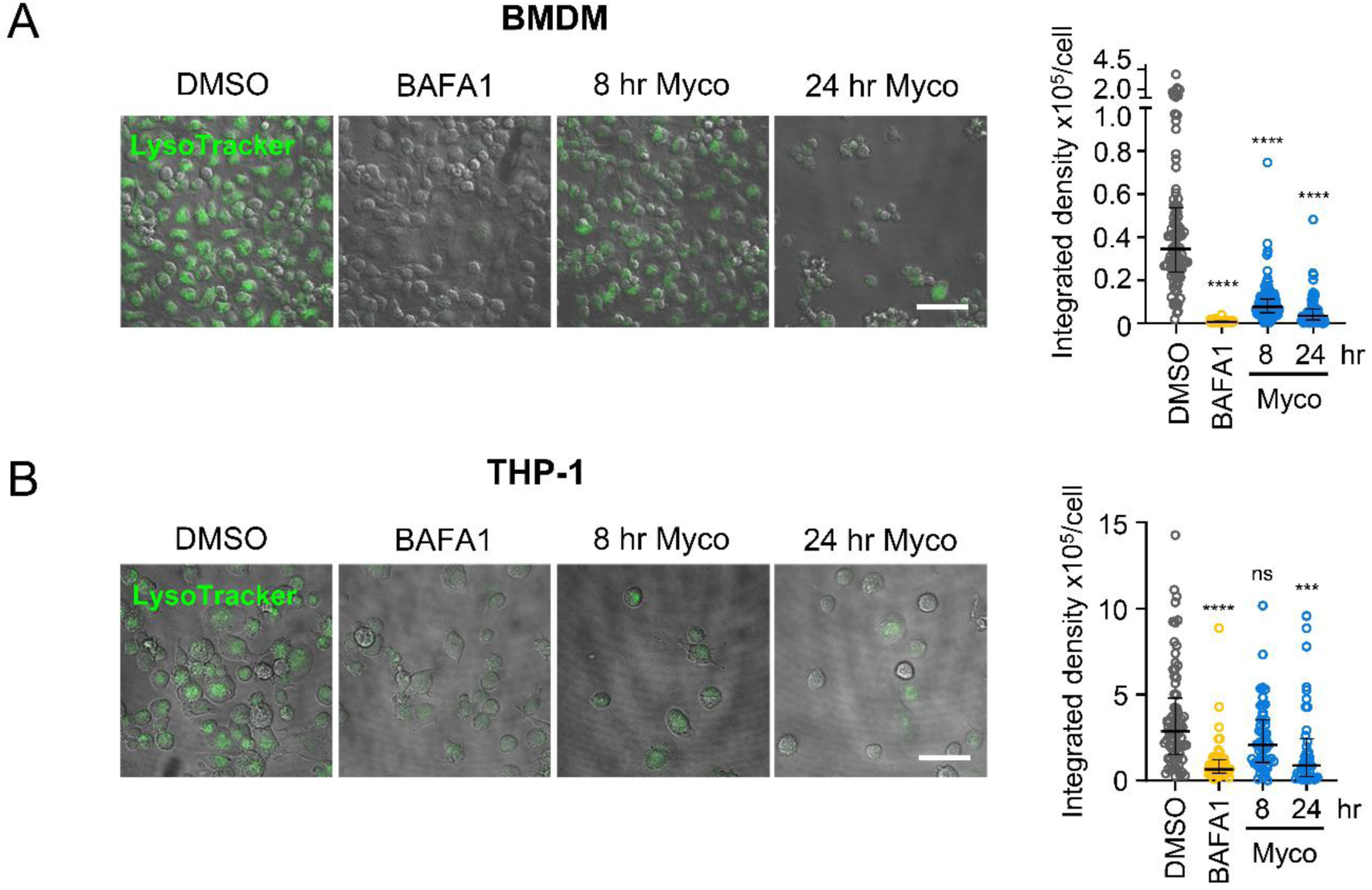
Prolonged exposure to mycolactone leads to a reduction in lysosome number and acidity. **A.** BMDM were seeded onto IBIDI 8-well chamberslides and incubated with 0.05% DMSO for 24hr, 31.25ng/ml mycolactone (Myco) for various times or 200nM Bafilomycin A1 (BAFA1) for 4hr, then incubated with 75nM Lysotracker Green DND-26 for 10min before washing. Cells were re-incubated 10min then imaged. Scale bar = 50µm. Data represents median of integrated density of Lysotracker fluorescence per cell for at least 74 cells ± IQR. Data representative of n=2 independent experiments. **B.** THP-1 cells were plated onto IBIDI 8-well chamber slides in the presence of 20ng/ml PMA. After 72hr medium was replaced and cells reincubated for 24hr before adding 0.05% DMSO for 24hr, 31.25ng/ml mycolactone (Myco) for various times or 200nM Bafilomycin A1 (BAFA1) for 4hr, then incubated with 75nM Lysotracker Green DND-26 for 10min before washing. Cells were re-incubated 10min then imaged. Scale bar = 50µm. Data represents median of integrated density of Lysotracker fluorescence per cell for at least 41 cells ± IQR. Data is representative of n=3 independent experiments. All statistical analysis was carried out using Kruskal-Wallis test. ns; not significant, ***; p<0.001, ****; p<0.0001.

## References

Abbas YM, Wu D, Bueler SA, Robinson CV, Rubinstein JL (2020) Structure of V-ATPase from the mammalian brain. Science 367: 1240–1246

Andrejewski N, Punnonen EL, Guhde G, Tanaka Y, Lullmann-Rauch R, Hartmann D, von Figura K, Saftig P (1999) Normal lysosomal morphology and function in LAMP-1-deficient mice. J Biol Chem 274: 12692–12701

Baron L, Paatero AO, Morel JD, Impens F, Guenin-Macé L, Saint-Auret S, Blanchard N, Dillmann R, Niang F, Pellegrini S et al (2016) Mycolactone subverts immunity by selectively blocking the Sec61 translocon. J Exp Med 213: 2885–2896

Bhadra P, Dos Santos S, Gamayun I, Pick T, Neumann C, Ogbechi J, Hall BS, Zimmermann R, Helms V, Simmonds RE et al (2021) Mycolactone enhances the Ca2+ leak from endoplasmic reticulum by trapping Sec61 translocons in a Ca2+ permeable state. Biochem J 478: 4005–4024

Bowman EJ, Siebers A, Altendorf K (1988) Bafilomycins: a class of inhibitors of membrane ATPases from microorganisms, animal cells, and plant cells. Proc Natl Acad Sci U S A 85: 7972–7976

Domenger A, Choisy C, Baron L, Mayau V, Perthame E, Deriano L, Arnulf B, Bories JC, Dadaglio G, Demangel C (2022) The Sec61 translocon is a therapeutic vulnerability in multiple myeloma. EMBO Mol Med 14: e14740

Feng R, Liu F, Li R, Zhou Z, Lin Z, Lin S, Deng S, Li Y, Nong B, Xia Y et al (2024) The rapid proximity labeling system PhastID identifies ATP6AP1 as an unconventional GEF for Rheb. Cell Res 34: 355–369

Gama JB, Ohlmeier S, Martins TG, Fraga AG, Sampaio-Marques B, Carvalho MA, Proenca F, Silva MT, Pedrosa J, Ludovico P (2014) Proteomic analysis of the action of the Mycobacterium ulcerans toxin mycolactone: targeting host cells cytoskeleton and collagen. PLoS Negl Trop Dis 8: e3066

George KM, Chatterjee D, Gunawardana G, Welty D, Hayman J, Lee R, Small PL (1999) Mycolactone: a polyketide toxin from Mycobacterium ulcerans required for virulence. Science 283: 854–857

George KM, Pascopella L, Welty DM, Small PL (2000) A Mycobacterium ulcerans toxin, mycolactone, causes apoptosis in guinea pig ulcers and tissue culture cells. Infect Immun 68: 877–883

Gerard SF, Hall BS, Zaki AM, Corfield KA, Mayerhofer PU, Costa C, Whelligan DK, Biggin PC, Simmonds RE, Higgins MK (2020) Structure of the Inhibited State of the Sec Translocon. Mol Cell 79: 406–415 e407

Guenin-Mace L, Carrette F, Asperti-Boursin F, Le Bon A, Caleechurn L, Di Bartolo V, Fontanet A, Bismuth G, Demangel C (2011) Mycolactone impairs T cell homing by suppressing microRNA control of L-selectin expression. Proc Natl Acad Sci U S A 108: 12833–12838

Guida MC, Hermle T, Graham LA, Hauser V, Ryan M, Stevens TH, Simons M (2018) ATP6AP2 functions as a V-ATPase assembly factor in the endoplasmic reticulum. Mol Biol Cell 29: 2156–2164

Hall BS, Dos Santos SJ, Hsieh LT, Manifava M, Ruf MT, Pluschke G, Ktistakis N, Simmonds RE (2021) Inhibition of the SEC61 translocon by mycolactone induces a protective autophagic response controlled by EIF2S1-dependent translation that does not require ULK1 activity. Autophagy: 1–19

Hall BS, Hill K, McKenna M, Ogbechi J, High S, Willis AE, Simmonds RE (2014) The pathogenic mechanism of the Mycobacterium ulcerans virulence factor, mycolactone, depends on blockade of protein translocation into the ER. PLoS Pathog 10: e1004061

Heaton NS, Moshkina N, Fenouil R, Gardner TJ, Aguirre S, Shah PS, Zhao N, Manganaro L, Hultquist JF, Noel J et al (2016) Targeting Viral Proteostasis Limits Influenza Virus, HIV, and Dengue Virus Infection. Immunity 44: 46–58

Hsieh LT, Hall BS, Newcombe J, Mendum TA, Varela SS, Umrania Y, Deery MJ, Shi WQ, Diaz-Delgado J, Salguero FJ et al (2025) The Mycobacterium ulcerans toxin mycolactone causes destructive Sec61-dependent loss of the endothelial glycocalyx and vessel basement membrane to drive skin necrosis. Elife 12

Itskanov S, Wang L, Junne T, Sherriff R, Xiao L, Blanchard N, Shi WQ, Forsyth C, Hoepfner D, Spiess M et al (2023) A common mechanism of Sec61 translocon inhibition by small molecules. Nat Chem Biol 19: 1063–1071

Jansen EJ, Timal S, Ryan M, Ashikov A, van Scherpenzeel M, Graham LA, Mandel H, Hoischen A, Iancu TC, Raymond K et al (2016) ATP6AP1 deficiency causes an immunodeficiency with hepatopathy, cognitive impairment and abnormal protein glycosylation. Nat Commun 7: 11600

Klionsky DJ, Abdel-Aziz AK, Abdelfatah S, Abdellatif M, Abdoli A, Abel S, Abeliovich H, Abildgaard MH, Abudu YP, Acevedo-Arozena A et al (2021) Guidelines for the use and interpretation of assays for monitoring autophagy (4th edition)(1). Autophagy 17: 1–382

Marion E, Song OR, Christophe T, Babonneau J, Fenistein D, Eyer J, Letournel F, Henrion D, Clere N, Paille V et al (2014) Mycobacterial toxin induces analgesia in buruli ulcer by targeting the angiotensin pathways. Cell 157: 1565–1576

Maxson ME, Abbas YM, Wu JZ, Plumb JD, Grinstein S, Rubinstein JL (2022) Detection and quantification of the vacuolar H+ATPase using the Legionella effector protein SidK. J Cell Biol 221

McKenna M, Simmonds RE, High S (2017) Mycolactone reveals the substrate-driven complexity of Sec61-dependent transmembrane protein biogenesis. J Cell Sci 130: 1307–1320

Nelson N, Harvey WR (1999) Vacuolar and plasma membrane proton-adenosinetriphosphatases. Physiol Rev 79: 361–385

Ogbechi J, Hall BS, Sbarrato T, Taunton J, Willis AE, Wek RC, Simmonds RE (2018) Inhibition of Sec61-dependent translocation by mycolactone uncouples the integrated stress response from ER stress, driving cytotoxicity via translational activation of ATF4. Cell Death Dis 9: 397

Ogbechi J, Ruf MT, Hall BS, Bodman-Smith K, Vogel M, Wu HL, Stainer A, Esmon CT, Ahnstrom J, Pluschke G et al (2015) Mycolactone-Dependent Depletion of Endothelial Cell Thrombomodulin Is Strongly Associated with Fibrin Deposition in Buruli Ulcer Lesions. PLoS Pathog 11: e1005011

Pankiv S, Clausen TH, Lamark T, Brech A, Bruun JA, Outzen H, Overvatn A, Bjorkoy G, Johansen T (2007) p62/SQSTM1 binds directly to Atg8/LC3 to facilitate degradation of ubiquitinated protein aggregates by autophagy. J Biol Chem 282: 24131–24145

Pareja F, Brandes AH, Basili T, Selenica P, Geyer FC, Fan D, Da Cruz Paula A, Kumar R, Brown DN, Gularte-Merida R et al (2018) Loss-of-function mutations in ATP6AP1 and ATP6AP2 in granular cell tumors. Nat Commun 9: 3533

Pena-Llopis S, Vega-Rubin-de-Celis S, Schwartz JC, Wolff NC, Tran TA, Zou L, Xie XJ, Corey DR, Brugarolas J (2011) Regulation of TFEB and V-ATPases by mTORC1. EMBO J 30: 3242–3258

Qi C, Lei L, Hu J, Wang G, Liu J, Ou S (2022) Identification of a five-gene signature deriving from the vacuolar ATPase (V-ATPase) sub-classifies gliomas and decides prognoses and immune microenvironment alterations. Cell Cycle 21: 1294–1315

Raben N, Puertollano R (2016) TFEB and TFE3: Linking Lysosomes to Cellular Adaptation to Stress. Annu Rev Cell Dev Biol 32: 255–278

Rehan S, Tranter D, Sharp PP, Craven GB, Lowe E, Anderl JL, Muchamuel T, Abrishami V, Kuivanen S, Wenzell NA et al (2023) Signal peptide mimicry primes Sec61 for client-selective inhibition. Nat Chem Biol 19: 1054–1062

Rujano MA, Cannata Serio M, Panasyuk G, Peanne R, Reunert J, Rymen D, Hauser V, Park JH, Freisinger P, Souche E et al (2017) Mutations in the X-linked ATP6AP2 cause a glycosylation disorder with autophagic defects. J Exp Med 214: 3707–3729

Sardiello M, Palmieri M, di Ronza A, Medina DL, Valenza M, Gennarino VA, Di Malta C, Donaudy F, Embrione V, Polishchuk RS et al (2009) A gene network regulating lysosomal biogenesis and function. Science 325: 473-477

Shi W, Lu D, Wu C, Li M, Ding Z, Li Y, Chen B, Lin X, Su W, Shao X et al (2021) Coibamide A kills cancer cells through inhibiting autophagy. Biochem Biophys Res Commun 547: 52–58

Stinear TP, Seemann T, Pidot S, Frigui W, Reysset G, Garnier T, Meurice G, Simon D, Bouchier C, Ma L et al (2007) Reductive evolution and niche adaptation inferred from the genome of Mycobacterium ulcerans, the causative agent of Buruli ulcer. Genome Res 17: 192–200

Torrado E, Fraga AG, Castro AG, Stragier P, Meyers WM, Portaels F, Silva MT, Pedrosa J (2007) Evidence for an intramacrophage growth phase of Mycobacterium ulcerans. Infect Immun 75: 977–987

Torrado E, Fraga AG, Logarinho E, Martins TG, Carmona JA, Gama JB, Carvalho MA, Proenca F, Castro AG, Pedrosa J (2010) IFN-gamma-dependent activation of macrophages during experimental infections by Mycobacterium ulcerans is impaired by the toxin mycolactone. J Immunol 184: 947–955

Tranter D, Paatero AO, Kawaguchi S, Kazemi S, Serrill JD, Kellosalo J, Vogel WK, Richter U, Mattos DR, Wan X et al (2020) Coibamide A Targets Sec61 to Prevent Biogenesis of Secretory and Membrane Proteins. ACS Chem Biol 15: 2125–2136

Wang L, Wu D, Robinson CV, Wu H, Fu TM (2020) Structures of a Complete Human V-ATPase Reveal Mechanisms of Its Assembly. Mol Cell 80: 501–511 e503

Xu L, Shen X, Bryan A, Banga S, Swanson MS, Luo ZQ (2010) Inhibition of host vacuolar H+-ATPase activity by a Legionella pneumophila effector. PLoS Pathog 6: e1000822

Yan Z, Huang A, Ma D, Hong C, Zhang S, He L, Rao H, Luo S (2025) ATP6AP1 promotes cell proliferation and tamoxifen resistance in luminal breast cancer by inducing autophagy. Cell Death Dis 16: 201

Yotsu RR, Suzuki K, Simmonds RE, Bedimo R, Ablordey A, Yeboah-Manu D, Phillips R, Asiedu K (2018) Buruli Ulcer: a Review of the Current Knowledge. Curr Trop Med Rep 5: 247–256

Zhang J, Zeng W, Han Y, Lee WR, Liou J, Jiang Y (2023) Lysosomal LAMP proteins regulate lysosomal pH by direct inhibition of the TMEM175 channel. Mol Cell 83: 2524–2539 e2527

